# A standardized framework for representation of ancestry data in genomics studies, with application to the NHGRI-EBI GWAS Catalog

**DOI:** 10.1101/129395

**Authors:** Joannella Morales, Emily H. Bowler, Annalisa Buniello, Maria Cerezo, Peggy Hall, Laura W. Harris, Emma Hastings, Heather A. Junkins, Cinzia Malangone, Aoife C. McMahon, Annalisa Milano, Danielle Welter, Tony Burdett, Fiona Cunningham, Paul Flicek, Helen Parkinson, Lucia A. Hindorff, Jacqueline A. L. MacArthur

## Abstract

**Background:** The accurate description of ancestry is essential to interpret and integrate human genomics data, and to ensure that advances in the field of genomics benefit individuals from all ancestral backgrounds. However, there are no established guidelines for the consistent, unambiguous and standardized description of ancestry. To fill this gap, we provide a framework, designed for the representation of ancestry in GWAS data, but with wider application to studies and resources involving human subjects.

**Result:** Here we describe our framework and its application to the representation of ancestry data in a widely-used publically available genomics resource, the NHGRI-EBI GWAS Catalog. We present the first analyses of GWAS data using our ancestry categories, demonstrating the validity of the framework to facilitate the tracking of ancestry in big data sets. We exhibit the broader relevance and integration potential of our method by its usage to describe the well-established HapMap and 1000 Genomes reference populations. Finally, to encourage adoption, we outline recommendations for authors to implement when describing samples.

**Conclusions:** While the known bias towards inclusion of European ancestry individuals in GWA studies persists, African and Hispanic or Latin American ancestry populations contribute a disproportionately high number of associations, suggesting that analyses including these groups may be more effective at identifying new associations. We believe the widespread adoption of our framework will increase standardization of ancestry data, thus enabling improved analysis, interpretation and integration of human genomics data and furthering our understanding of disease.

## Background

The past 15 years have seen a dramatic growth in the field of genomics, with numerous efforts focused on understanding the etiology of common human disease and translating this to advances in the clinic. Genome-wide association studies (GWAS), in particular, are now a well-established mechanism to identify links between genetic variation and human disease[1–3]. The NHGRI-EBI GWAS Catalog[2,4], one of the largest repositories of summary GWAS data, contains over 3,000 publications and 43,000 SNP-trait associations as of July 2017. The Catalog is indispensable for researching existing findings on common diseases, enabling further investigations to identify causal variants, understand disease mechanisms and establish targets for treatment[5–8].

Essential to the interpretation and application of genomic data is the accurate description of the ancestry of the samples studied. Levels of genetic diversity and patterns of linkage disequilibrium (LD) vary by ancestry, with important implications for the experimental design, the generalizability of results and the identification of causal variants. Standardized ancestry representation is necessary to integrate data from different sources for further analysis and to enable robust search functionalities in bioinformatics resources. There are currently no established guidelines for the characterization and classification of ancestral background information. This has led to ambiguity and inconsistency, along with challenges in accessing and integrating data.

The need for genetic studies in more ancestrally diverse populations has been repeatedly articulated[9], most recently by Popejoy and Fullerton[10] and by Manolio, et al.[11]. The benefit of including diverse populations extends throughout the translational research spectrum, from GWAS discovery efforts to genomic medicine, for which variant interpretation can be greatly aided by ancestrally diverse sequence information[12,13]. Although inclusion efforts are improving over time, it is challenging to assess the status of such efforts, or to implement improved approaches, without a standardized way of representing ancestry data.

To fill these gaps, we here provide a framework to systematically describe and represent detailed ancestry information. Our method is driven by an analysis of data we manually curated from over 3,000 GWAS publications. We developed it for immediate application to the GWAS Catalog to make curated data accessible, searchable and compatible with other genomics data. However, the framework is broadly applicable to studies and resources involving human subjects, and its widespread adoption will enable improved analysis, interpretation and integration of data, ultimately furthering our understanding of the genetic architecture of disease in different human populations.

## Results

### Standardized ancestry framework

We developed the framework to enable the generation of comprehensive and standardized representation of ancestry information for samples included in GWAS. We had several motivations: 1) our observation, from curating thousands of publications, that ancestry data is often poorly represented, inconsistent or even completely absent, 2) the requirement to describe, represent and store ancestry data in a manner that allows robust searching and visualization of data in the GWAS Catalog, 3) the increased interest in ancestry from the scientific community and 4) the need to analyze the ancestry of samples, assess diversity and generate metrics that would allow the community to identify and address gaps in this area.

Our framework involves representing ancestry data in two forms: (1) a detailed sample description and (2) an ancestry category from a controlled list (Table 1). Detailed descriptions aim to capture accurate, informative and comprehensive information regarding the ancestry or genealogy of the samples. Ancestry categories are used to establish hierarchical relationships between groups and populations. We believe this dual representation of ancestry is both informative and useful. The two types of descriptions complement each other. The detailed description is granular and heterogeneous, whereas the use of a limited number of categories reduces complexity, facilitating data representation, integration, searchability and further analyses. The framework also allows for country information to be recorded providing additional detail on sample demographics.

**Table 1.**
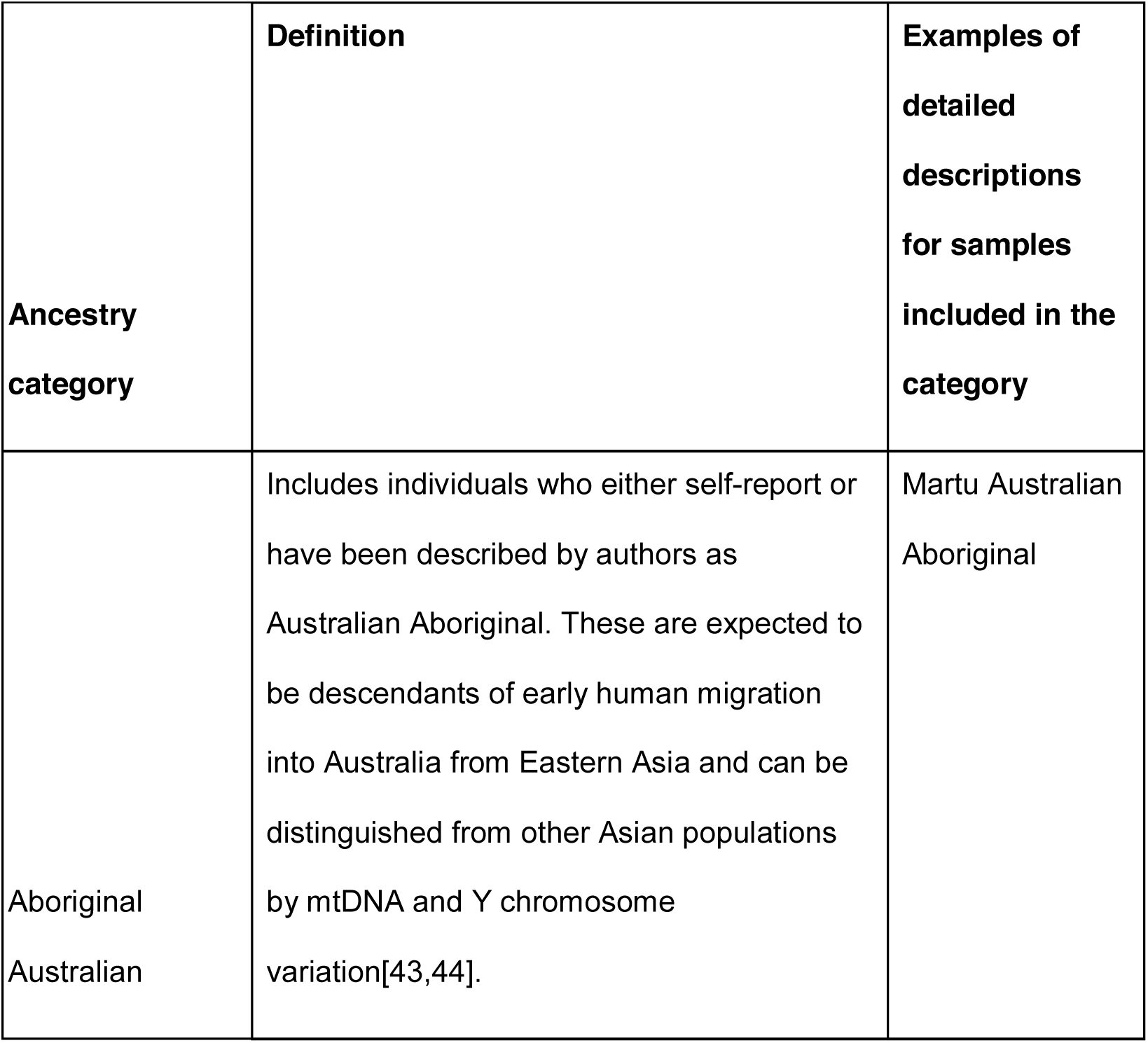
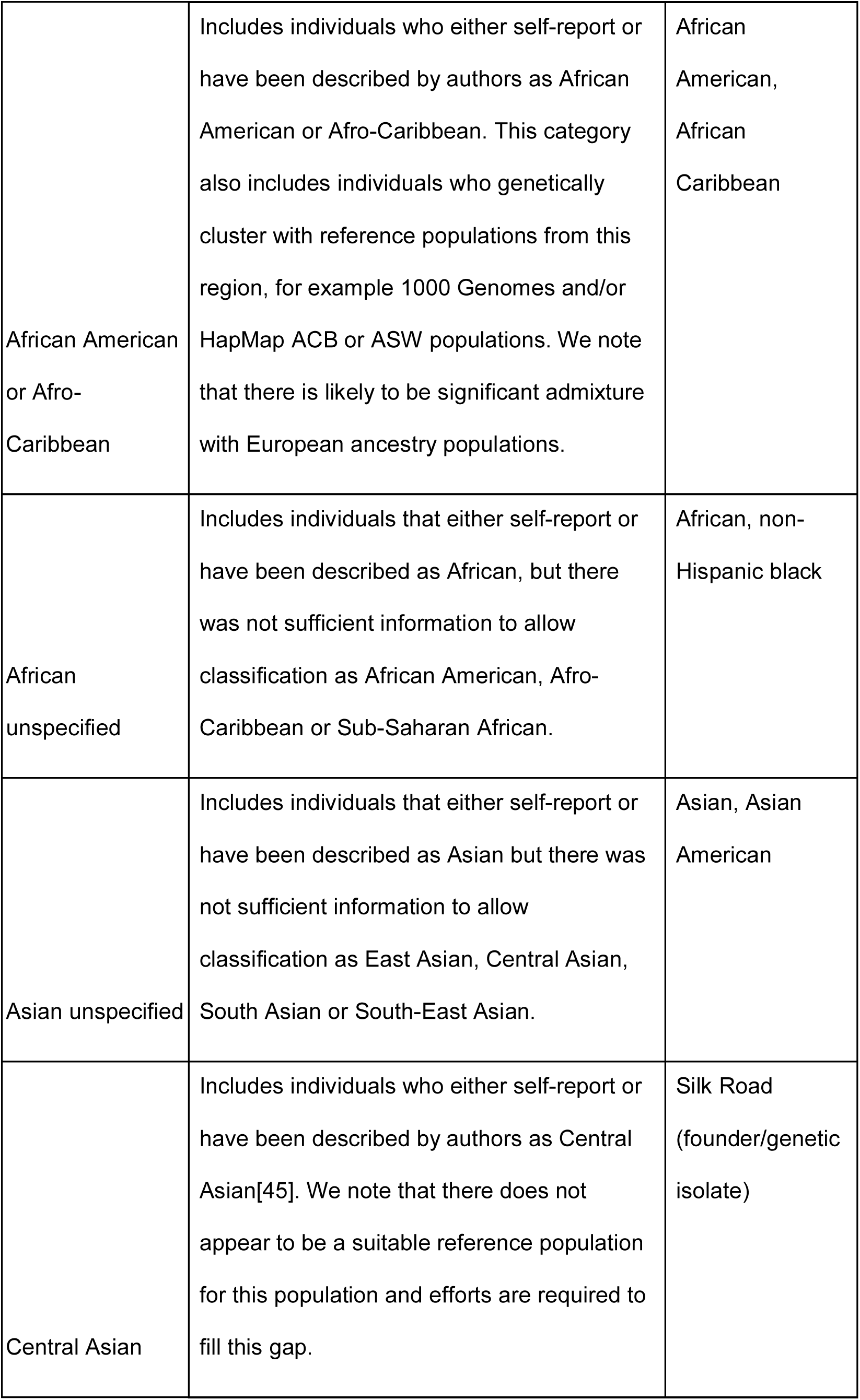
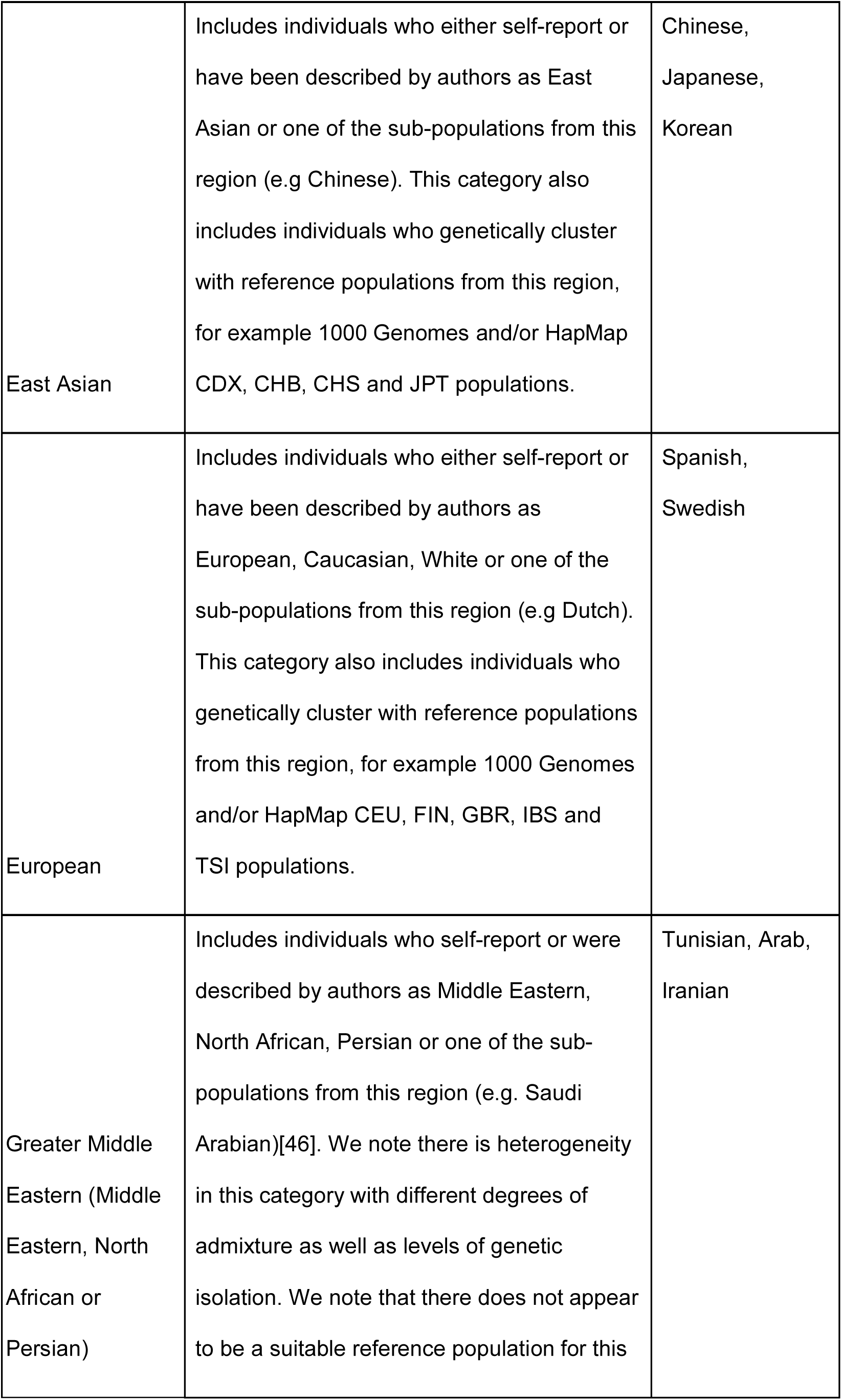
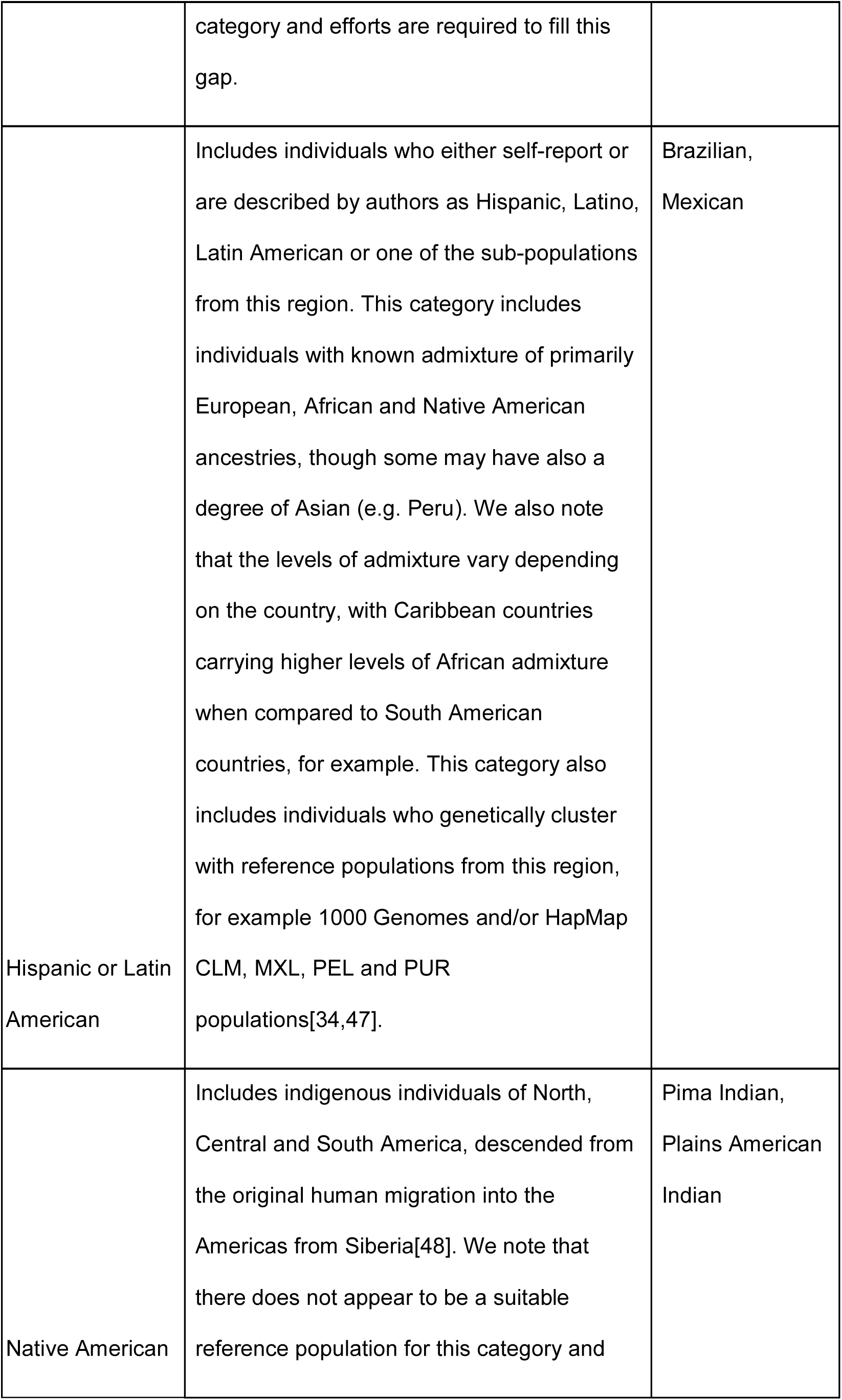
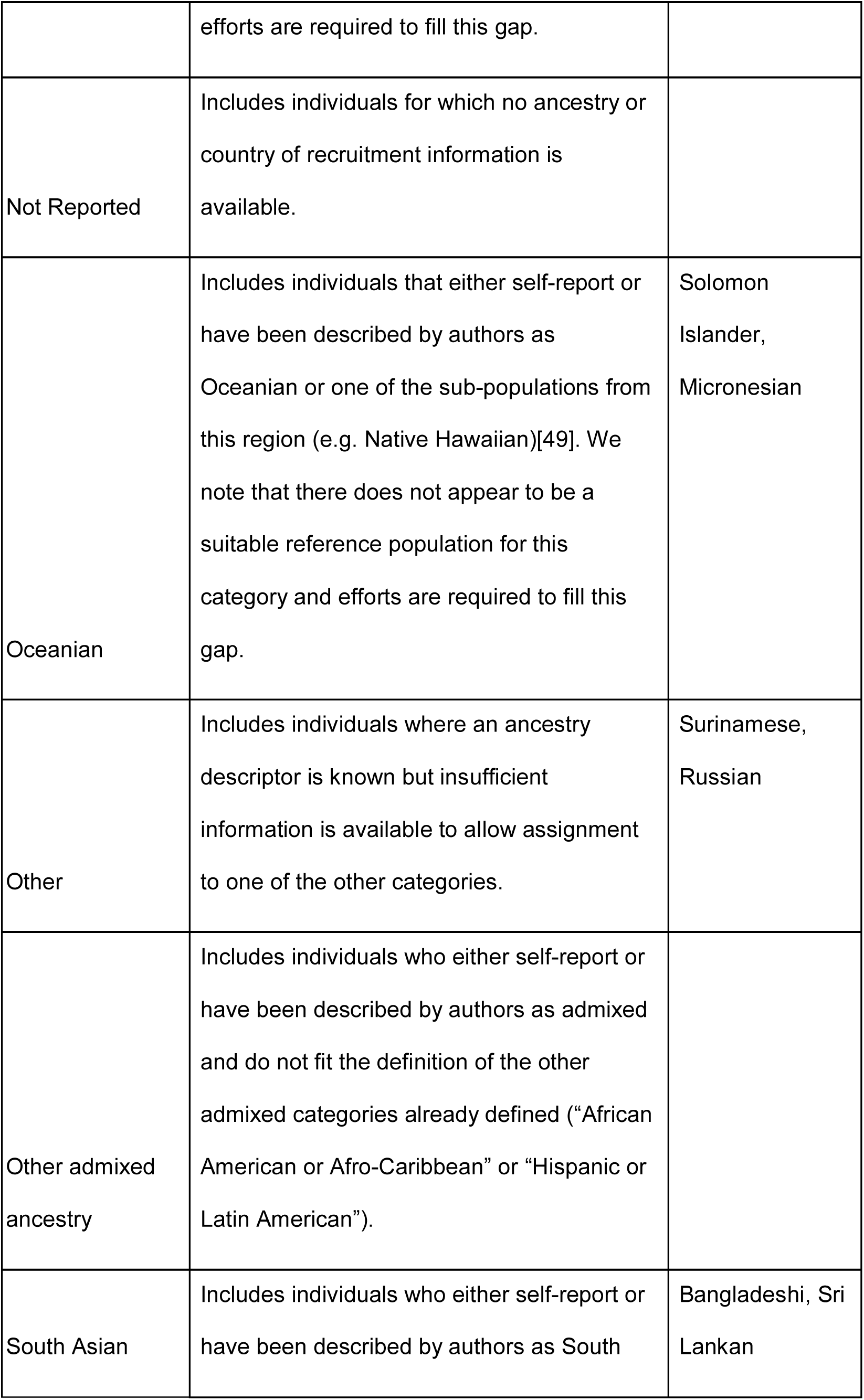
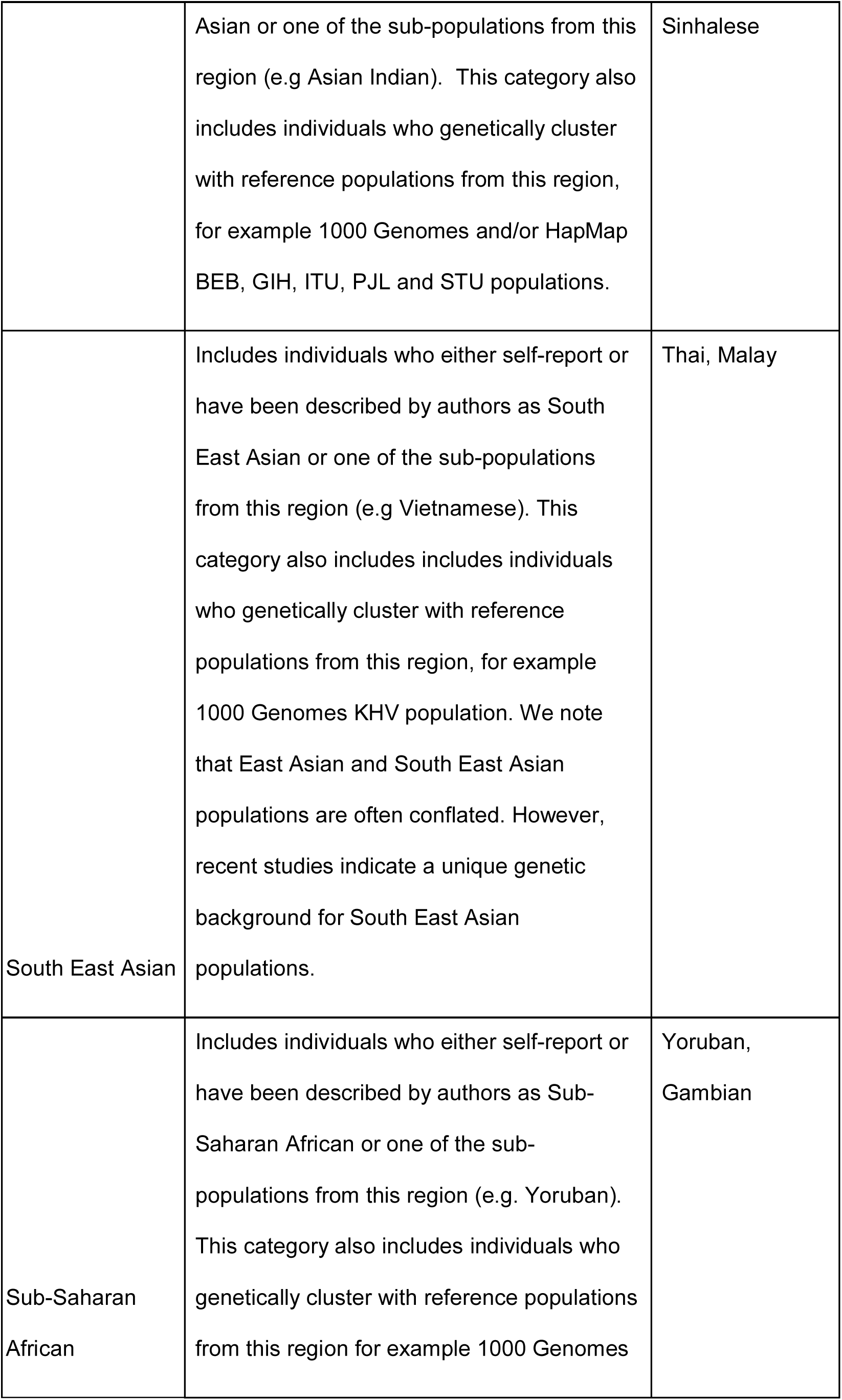
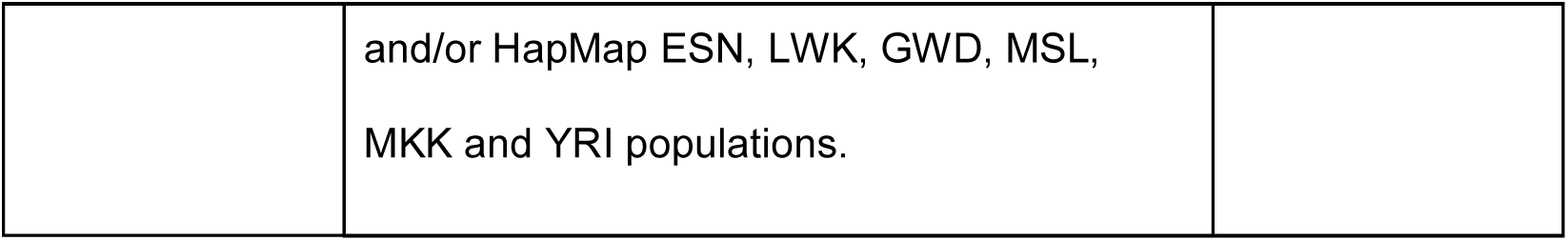
Ancestry categories. Distinct regional population groupings used in this framework. They are assigned to cohorts with distinct and well-defined patterns of genetic variation, in addition to individuals with inferred relatedness to these cohorts. A full list of GWAS Catalog sample descriptions assigned to each category can be found in supplementary table 2.

### Application of the framework to the GWAS Catalog and other resources

Application of this framework by curators to samples reported in publications relies on manual interpretation and extraction of author-reported data. To ensure consistent application by GWAS Catalog curators, we created a set of comprehensive data extraction guidelines (Supplementary Note). When the information provided by authors is limited or ambiguous, we consider country of recruitment demographics and peer-reviewed population genetics publications.

We have now generated detailed descriptions and assigned ancestry categories to 3,000 publications, representing 4,000 separate GWA studies and 83 million individuals, as of July 2017. A full list of all detailed descriptions currently included in the Catalog is provided in Supplementary Table 1. Examples that illustrate how the framework was applied to Catalog samples can be found in Supplementary Table 2. All curated ancestry data is available from the GWAS Catalog website[4] (Figure 1) and via download[14].

**Figure 1.**
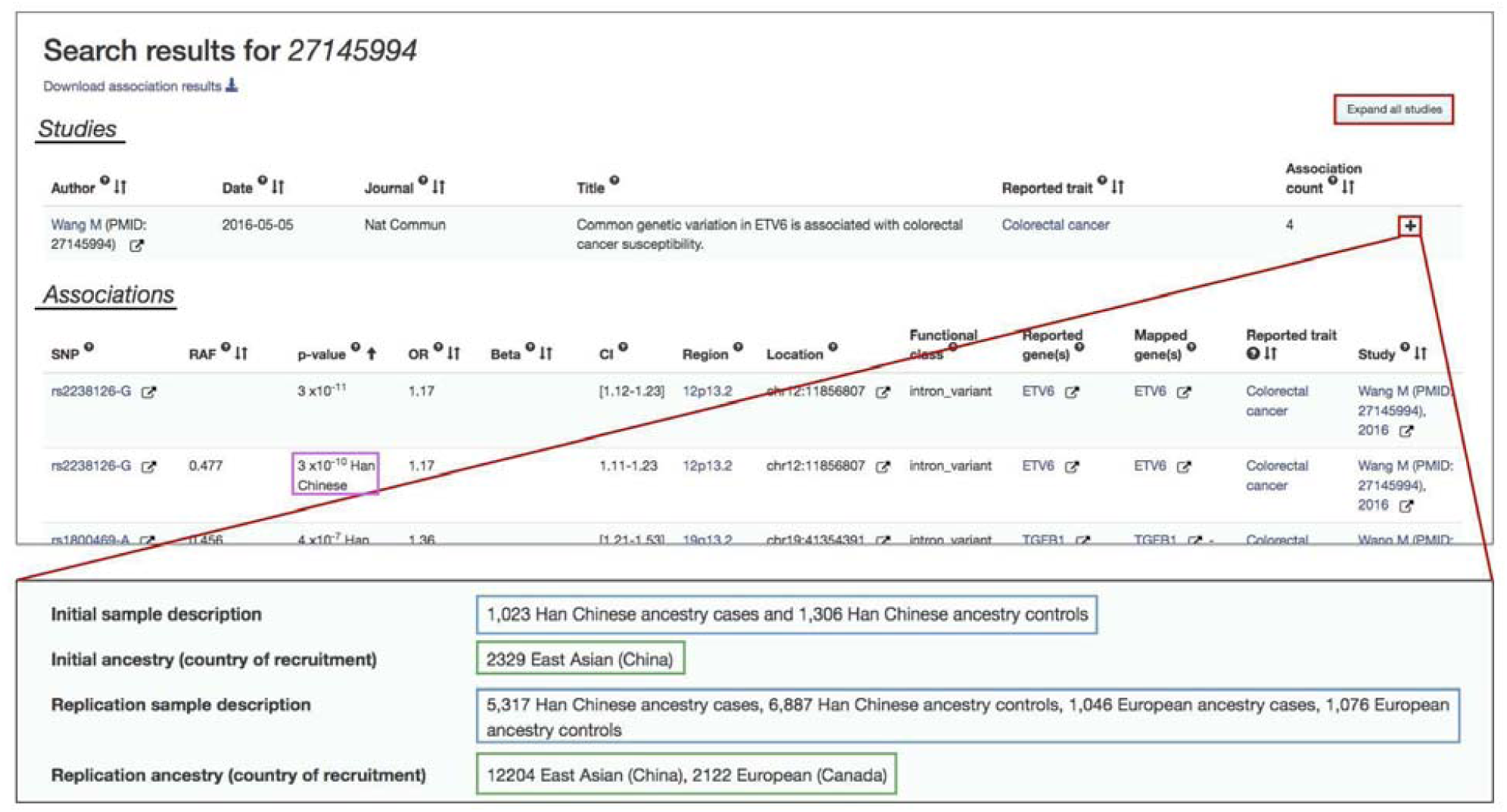
Representation of ancestry data in the GWAS Catalog search interface (www.ebi.ac.uk/gwas). Ancestry-related data is found in the Studies and Associations tables (underlined in black) when searching the Catalog. This figure shows the results of a search for PubMed Identifier 27145994. The sample description can be found in the Studies table, either by pressing “Expand all Studies” or the “+” on the study of interest (highlighted in red). Sample ancestry is captured in 2 forms: (1) Detailed description (highlighted in blue) and (2) Ancestry category (highlighted in green). The latter follows the format: sample size, category, (country of recruitment). In cases where multiple ancestries are included in a study, the ancestry associated with a particular association is found as an annotation in the p-value column in the Associations table (highlighted in pink).

#### Detailed description

The detailed description aims to accurately represent the ancestry or genealogy of each distinct group analyzed in a specific study in detail, as reported by the author. Information about the homogeneity of the samples, including whether the cohort is admixed or taken from a founder or isolated population, is included. In the GWAS Catalog, the majority of the detailed descriptions include terms that describe the location of participants’ ancestors over the past few generations (“French”, “Japanese”), while admixed populations are primarily described using ethnic descriptors (“Hispanic”). Isolated populations are described using either location or ethnicity terms in addition to being described explicitly as genetically isolated (“Old Order Amish (founder or genetic isolate) population”, “Norfolk Island (founder or genetic isolate) population”.

#### Ancestry categories

Ancestry category assignment from the list presented in Table 1 requires careful consideration. When clearly stated, author-reported categories are extracted, with precedence given to genetically-inferred data. If a category is not stated, curators infer the category based on the detailed description for the sample, which, as noted above, represents author-provided information.

In the absence of any ancestry data, the category “Not Reported” is assigned, unless geographical location of sample recruitment is stated. In such instances, curators infer ancestry from external sources, such as the United Nations[15] and The World Factbook[16]. Selecting a category for samples that derive from a country with a homogenous demographic composition, such as Japan, is straightforward. However, for samples from populations with limited known genetic genealogy, such as Azerbaijan, or for samples recruited in countries with ancestral diversity, such as Singapore, assigning a category is more challenging. These sources are particularly useful to obtain geographical and country-specific population information. The World Factbook is a regularly updated, comprehensive compendium of worldwide demographic data, covering all countries and territories of the world. However, since it does not necessarily provide ancestry data, the World Factbook is consulted when the only known information is the country of recruitment of samples. We expect that as increased care is taken to accurately report ancestry data, reliance on this resource will decrease. Peer-reviewed population genetic studies that characterize the genetic background of a given population may also be consulted. This is particularly helpful in cases where the sample cohort self-reported or is described using geographical or ethno-cultural terms, such as “Scandinavian” or “Punjabi Sikh”. Supplementary Table 3 provides a list of countries for which external sources were consulted. If the ancestry data provided in publications does not allow the resolution of samples into ancestrally distinct sets, more than one category may be selected from the list in Table 1, such as in the Catalog entry for Jiang R *et. al.* [17][18].

#### Country information

Country of recruitment (Figure 1) and country of origin provides additional demographic information and is extracted for each distinct sample set. Country of origin or recruitment is author-reported and not inferred from ancestry data. An exception is made for occasions when authors combine country of recruitment with an ancestry description (“Singaporean Chinese”). In these cases, we infer the country of recruitment (“Singapore”) although it is not explicitly stated. Country of origin is defined as the country of origin of the study participant’s grandparents or as the genealogy of the participants dating several generations.

### Wider application of the framework

The HapMap[19] and 1000 Genomes[20] projects have delivered a comprehensive survey of human genetic variation across worldwide populations. The application of our method to the ancestry representation of these reference populations therefore provides huge integration potential and demonstrates the relevance of our framework beyond GWAS publications.

For all populations, we assigned ancestry category, country of recruitment, country of origin and a detailed description, if provided by each project (Supplementary table 4). We are developing an integrated tool to provide access to population genetic and linkage disequilibrium data from these reference populations. This will be available in the near future.

To facilitate the application of our approach to databases and resources, we have developed an ancestry-specific ontology based on our framework. We have defined terms, identified synonyms and established hierarchical relationships between all curated terms and categories. Integration of the ontology into any search interface will enable users to perform more powerful and precise ancestry-related queries[21]. We aim to integrate it into the GWAS Catalog website in the near future. The ancestry ontology[22] can be browsed and downloaded (manuscript in preparation).

### Application of framework to assess changes in diversity

Several members of the community have called for greater efforts to increase diversity in genomics studies[10]. However, it is difficult to assess progress or identify concrete areas for improvement without a method to easily track changes in ancestry data. Our framework addresses this challenge by requiring the use of specific categories when describing ancestry. This process establishes hierarchical relationships between populations, thus facilitating reproducible tracking of changes in standardized categories over time. A number of authors[9,10] have reviewed the ancestry distribution in the Catalog, but focused exclusively on the detailed descriptions, which are heterogeneous as they are based on the authors’ language. Here we present the first analyses using our ancestry categories and demonstrate the validity of our framework to facilitate the tracking of ancestry in big data sets.

As expected, similar to previous reports[10], we found that the majority (78%) of individuals in the Catalog are exclusively of European ancestry (Figure 2a). The second largest group includes individuals of Asian descent (11%), with East Asians comprising 9% of the Catalog’s samples. The disproportionate focus on Europeans was more prevalent in the earlier years of the Catalog (86% of individuals in studies published between 2005 and 2010; 76% between 2011 and 2016, Figure 3). The reduced proportion of European ancestry individuals added to the Catalog in the last 5 years reflects an increase in Asian (7.5% to 11.4%, 1.5-fold increase), African (0.8% to 2.8%, 3.5-fold increase), Hispanic or Latin American (0.1% to 1.2%, 9-fold increase) and Middle Eastern (0.01% to 0.08%, 7-fold increase) samples. Though the proportion of Hispanic or Latin Americans exhibited the largest increase, when considering the absolute number of individuals, the largest increase, by far, came from Asian populations; Asian ancestry individuals increased from almost 900,000 in the first 5 years to over 7 million added to the Catalog in the last 5 years, compared to an increase of 721,000 Hispanic or Latin Americans.

**Figure 2.**
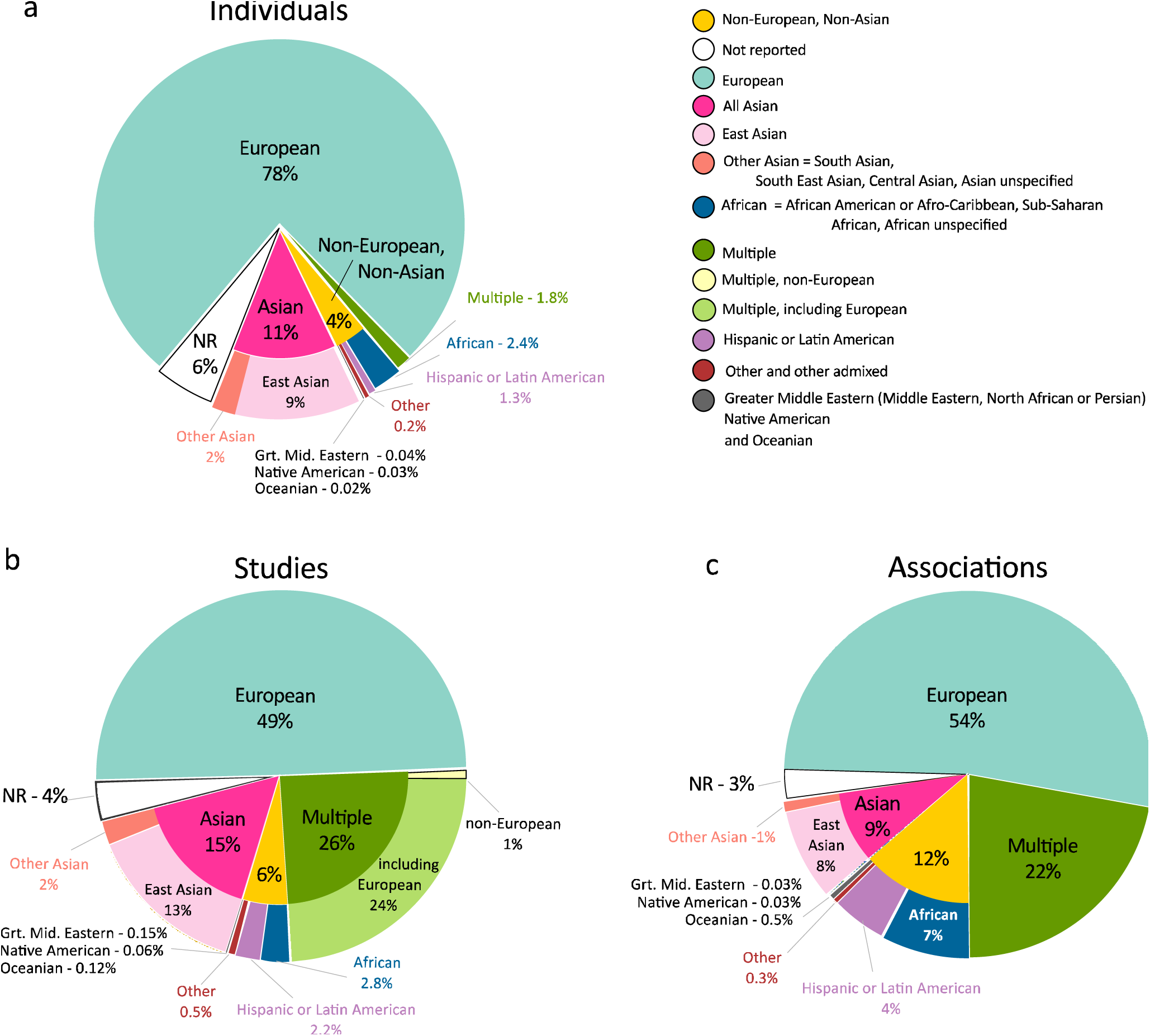
Ancestry category distribution in the GWAS Catalog. This figure summarizes the distribution of ancestry categories in percentages, of individuals (N=83,200,468; panel a), studies (N= 4,100; panel b) and associations (N=43,919; panel c). The largest category in all panels is European (aqua). At the level of individuals (a), the largest non-European category is Asian (bright pink), with East Asian (light pink) accounting for the majority. Non-European, Non-Asian categories together (yellow) comprise 4% of individuals, and there are 6% (white) of samples for which an ancestry category could not be specified. Panel c demonstrates the disproportionate contribution of associations from African (blue) and Hispanic/Latin American (purple) categories, when compared to the percentage of individuals (a, blue, purple, respectively) and studies (b, blue, purple, respectively).

**Figure 3.**
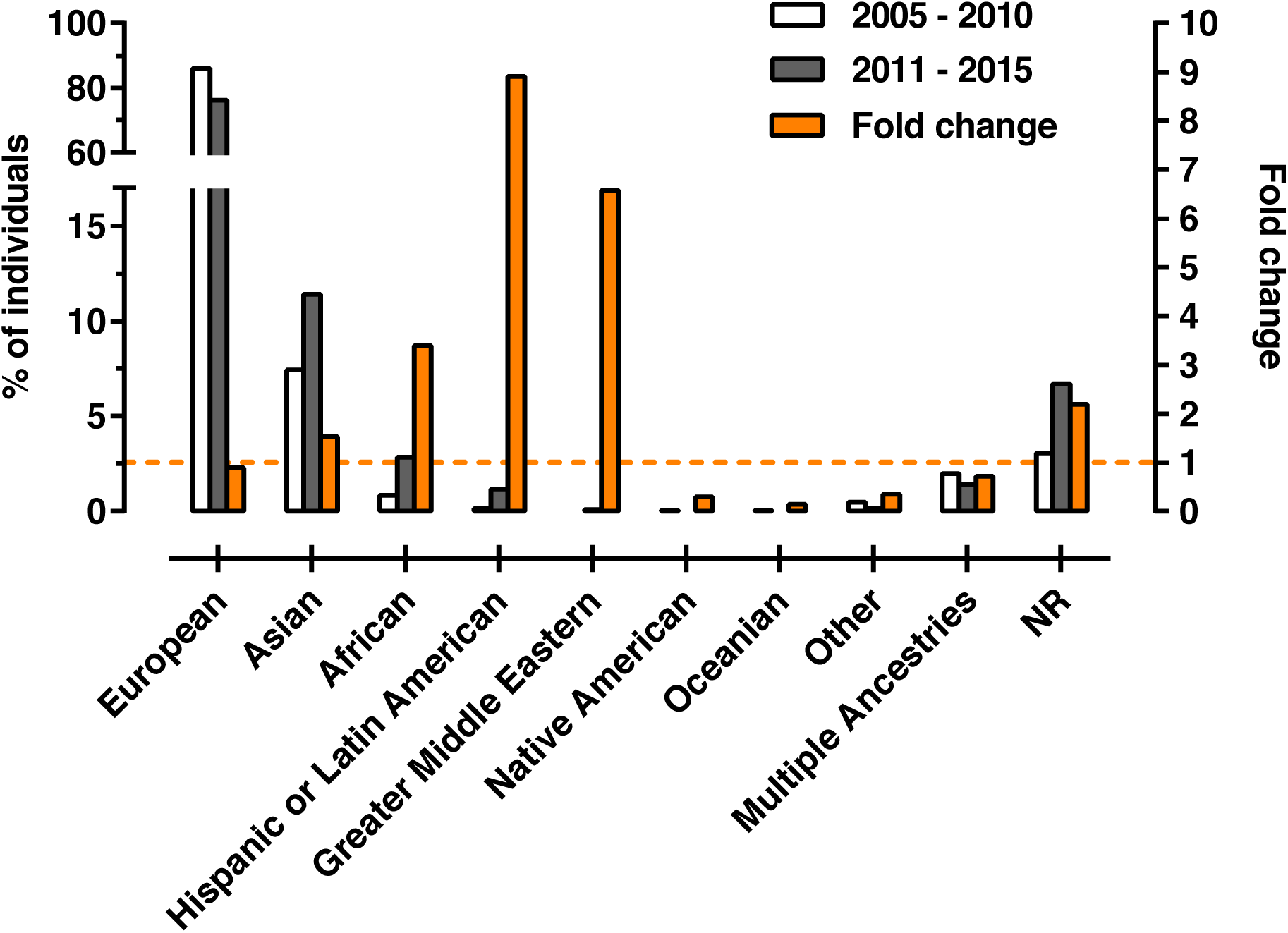
Distribution of individuals in GWAS Catalog studies published between 2005 – 2010 compared to 2011 – 2016. This figure displays the distribution of individuals in percentages, included in the 915 studies published between 2005 – 2010 compared to the distribution of individuals included in the 2,905 studies published between 2011 – 2016.

A similar trend is observed when analyzing the ancestry distribution in independent GWA studies. Approximately 50% of all studies are performed on exclusively European ancestry individuals, with an additional 24% of studies including some samples of European descent (Figure 2b). A more granular analysis of the traits with the largest number of GWAS in the Catalog presented the same European bias, with 57-80% of studies, depending on the trait, carried out in European ancestry individuals, followed by East Asians (7-28% of studies) (Supplementary Fig. 3). In studies that analyzed multiple ancestries, the vast majority (> 90%) include some European ancestry individuals, regardless of the trait. The traits that display the largest proportion of ancestral diversity are anthropometric traits, such as body mass index (BMI) and height, and common diseases, including type 2 diabetes and cardiovascular disease (Supplementary Fig. 3).

Interestingly, when we focused our analysis on the number of associations identified in each ancestry category, we noted a different distribution to the ancestry distribution of individuals (Figure 2c). This disparity is particularly pronounced for studies including African or Hispanic or Latin American samples; African ancestries contribute 2.4% of individuals but 7% of associations, while Hispanic/Latin Americans contribute 1.3% of individuals compared to 4.3% of associations. The opposite effect was seen in Europeans, with 54% of associations compared to 78% of individuals. In addition, we also observed a disproportionate number of associations contributed by the “Multiple ancestries” category, likely reflecting the Catalog’s inclusion of trans-ethnic meta-analyses and replication efforts in diverse ancestries.

### Recommendations to authors

The analysis of the over 3,000 GWAS publications revealed inconsistent and ambiguous reporting of ancestry data, with a significant percentage of studies (∼ 4%) not reporting any ancestry information at all. Given that there are no established guidelines for the description of ancestry, and in an effort to assist the community as it seeks to improve in this area, we here provide a set of specific recommendations for authors, also summarized in Box 1. We believe implementation of these recommendations will improve the quality of reporting and have a positive impact on the interpretation of published results, data reuse and reproducibility.

We recommend that authors make every effort to generate a detailed description for each distinct set of individuals included in their studies. Authors should also assess whether the genetic diversity of each distinct set is representative of one of the known populations listed and defined in Table 1, and indicate the corresponding category in the publication. If authors have no knowledge about the ancestry of the participants, are not able to determine it or cannot share it due to confidentiality concerns, we suggest noting this explicitly in the publication.

Where possible authors should provide genetically inferred, in addition to self-reported, ancestry information as the latter is often not an accurate representation of the underlying genetic background. Software to assess and control for ancestry is readily available and computationally feasible[23–28]. Encouragingly the proportion of studies that assess ancestry by genetic methods has increased, from 25% in the early days of GWAS (the first 100 eligible publications, 2005-2008) to 57% in 2016 (the first 100 eligible publications; Supplementary Fig. 4). Thus, we suggest authors consider using the methods listed in Supplementary Box 1 when inferring the ancestry of samples.

In general, terms that pertain to an individual’s ethno-cultural background should be avoided, unless this provides additional information regarding the genealogy of the samples. In such cases a descriptor that accurately reflects the underlying genetics should also be provided. For example, when describing “Punjabi Sikh” participants, every effort should be made to assess the genetic background and to indicate this in the publication, for example by stating “Punjabi Sikh South Asian ancestry individuals” rather than simply “Punjabi Sikh” or “Sikh”.

Particular care should be taken to note if a sample derives from founder or genetically isolated population; given their homogeneity and reduced genetic variation, these populations are especially well-suited for GWAS[29] and are increasingly used as sample sources. To reduce ambiguity, when describing isolates, the broader genetic background within which the population clusters should also be indicated. For example, Old Order Amish participants should be described as “Old Order Amish population isolate individuals of European ancestry”. While describing admixed populations can be challenging due to varying levels of admixture, every effort should be made to explicitly note whether the population is admixed and the ancestral backgrounds that contribute to admixture. For example “Hispanics/Latinos are ethnically heterogeneous, with admixture of European, West African, and Amerindian ancestral populations”, as stated in Hodonsky 2017[30].

## Discussion

### Summary

In this article we describe a framework for the standardized representation of ancestry data from genomics studies. Our method provides structure to unstructured data, enabling robust searching across large datasets and integration across resources. We have established validity by application to the over 3,000 GWAS publications currently in the GWAS Catalog and performed a detailed analysis. These data represent over 83 million individuals from diverse ancestral backgrounds, and, as such, it is likely one of the largest and most widely-used repositories of curated ancestry data. We have demonstrated relevance to, and integration potential with, other data types and studies by using our method to describe well-characterized reference populations, such as the HapMap and 1000 Genomes project populations. This will greatly facilitate integration of studies involving these populations with data included in the Catalog, and, indeed, with any other resource that implements our framework. We display the utility of the ancestry categories to simplify the tracking of efforts towards diversity, allowing the identification of gaps and highlighting specific areas for improvement. Interestingly, in addition to confirming known biases, our category-based analyses revealed that African and Hispanic or Latin American ancestry populations contribute a disproportionately high number of associations, suggesting that analyses including these groups may be more effective at identifying new associations. Finally, stemming from our extensive manual review of publications, we note a lack of current standards with regard to ancestry reporting and offer recommendations to authors to implement when describing their samples. This, we believe, will increase consistency and reduce ambiguity, facilitating the interpretation of results.

### Limitations to the framework

There are challenges inherent to both the design of the framework and its application. We recognize the sensitivities surrounding the concepts of race, ethnicity and ancestry, and that these terms are often used interchangeably without making a distinction between physical appearance, cultural traditions and genetic variation. This conflation can often be observed in censuses and other demographic tools, influencing how individuals and communities describe their background. The United States Census, for example, defines “White” as “a person having origins in any of the original peoples of Europe, the Middle East, or North Africa”[31], even though Middle Eastern and North African populations are known to cluster, in genetic analyses, independently from European ancestry populations. We thus recommend that authors continue to move away from relying solely on self-reported information and, as much as possible, also use genomic mechanisms to infer and describe the ancestry of participants.

We are aware that classifying the global human population is a challenging endeavor and the subject of numerous publications by expert population geneticists. Due to human evolution and patterns of migration, the ancestry of a particular population is complex and its definition is dependent on time. It is generally accepted that all modern human populations originated in Africa, and, due to the relatively small amount of genetic variation between populations from disparate geographical locations[20], genetic diversity among modern populations may be more suitably described as a continuum. However, it is both possible and useful to generate informative groupings. Reference populations, such as those included in the HapMap and 1000 Genomes Projects, or ancestry informative markers[32] that allow populations to be distinguished, have been characterized, and methods have been developed to adjust for population stratification and separate samples into clusters. In fact, these analyses between and within populations have demonstrated that clusters identified through genomic methods generally align with geographical and regional groupings [33]. Taking this into account, we designed our framework such that our categories closely resemble classifications currently in use by the community and defined by experts.

We do not view our categories as exhaustive or static. We envision that as more cohorts from diverse populations are characterized, there might arise a need to create additional categories or sub-categories. In addition, anticipating that admixture is likely to increase in the future, due to migration, for example, we also created categories to represent known (for example, “Hispanic or Latin American”) and emerging (for example, “Other admixed ancestries”) admixed groups. We recognize that classification of admixed samples is particularly challenging. The degree and type of admixture may vary within the population, and the accuracy of classification requires well-defined reference samples, which are lacking for some groups. As the community moves towards genetically-inferred ancestry descriptions, our categories are likely to become more precise and granular over time.

We believe there are benefits to utilizing categories. Practically, categories facilitate data integration and allow robust searches, which is an essential component of databases such as the GWAS Catalog. Also, the use of categories can be useful when carrying out further analyses. For example, querying the generalizability of results, identifying ancestry-specific associations or utilizing linkage disequilibrium information from a reference population to identify independent signals. We did not set out to define novel or authoritative global ancestral classifications. Rather, we aim to formalize and encourage the use of existing classifications to increase standardization and improve resource functionality, ultimately enabling more robust scientific analyses.

### Assessing diversity in genomics

Several reports have been published urging the scientific community to ensure that individuals from all ancestry backgrounds benefit from advances in the field of genomics[9,10]. Our proposed framework lays out a mechanism for the generation of consistent and comprehensive ancestry descriptions and this, in turn, facilitates the establishment of metrics and ultimately, the tracking of ancestry data over time.

Our analysis of individuals in GWA studies, using ancestry categories rather than Catalog detailed descriptions, as carried out in previous studies, confirms the persistent bias towards inclusion of European ancestry samples and the modest trend towards increased diversity. Our more robust analysis of approximately 43,000 associations found a disproportionately larger proportion of associations derived from African and Hispanic or Latin American populations, many of which have significant African admixture[34], than is expected based on the proportion of individuals. We suggest that the higher degree of genetic diversity and reduced linkage disequilibrium (LD) in African populations[35] offers an explanation for this result. Shorter LD blocks in African populations facilitate the separation of nearby but independent signals in a way that is more challenging in European populations, in which LD blocks tend to be longer. Additionally, the inclusion of larger numbers of individuals from African or Hispanic or Latin American populations allows for the identification of variants specific to these ancestries. Together, these observations suggest that utilizing samples from diverse populations for genomic studies may be advantageous and yield increased and more comprehensive results.

There are limitations to our analysis. First, considering that some cohorts have been included in numerous GWAS, it is highly likely that some individuals are represented multiple times in the Catalog. The impact of this is the skewing of results towards commonly-used or publicly available cohorts, which are perhaps likely to be of European or Asian ancestry. Another limitation stems from our criteria for inclusion of associations. Since we only include variants with a p-value < 1x10^-5^ and only the “index” variant at each locus, our analysis does not take into account all associations. To address this and make the Catalog more comprehensive, we now also include in the Catalog published summary statistics[36]. Finally, we were unable to assign a category to associations identified in studies that include multiple ancestries. This may be a factor contributing to the reduced number of associations derived from European populations, since the vast majority of multiple ancestry studies include Europeans (Figure 2b).

The analysis of diverse ancestries is advantageous from a scientific perspective. No one population contains all human variants[37], and alleles that are rare in one population may be common in a different population and thus easier to detect. Studies of diverse populations may also aid in fine mapping of existing signals or in identifying population-specific functional variation[37,38]. Variant interpretation for genomic medicine in ancestrally diverse or admixed populations relies on the availability of non-European allele frequencies, with potentially serious clinical consequences if such data are not available[12]. Finally, disease burden of common or complex diseases (e.g., cardiovascular disease or cancer) disproportionately impacts non-European populations. Of the commonly studied traits, the largest diversity of backgrounds was found for common anthropometric traits, heart disease, and type 2 diabetes. This is perhaps not surprising considering that metrics for these traits are easy to obtain, and the two diseases are among the top ten causes of death around the world, according to the World Health Organization[39]. It is also consistent with the observation that diseases for which global disease burden is substantial tend to lead to increased funding and research infrastructure. While we are encouraged by the trend we have seen in recent years towards increased diversity, we note that there are still very clear gaps as some groups continue to be underserved or ignored. We strongly urge the scientific community to expand their efforts to assemble and analyze cohorts, including especially underrepresented communities.

### Recommendations to authors

Our analysis also validates the need for a framework to improve the description of ancestry. Approximately 5.8% of individuals in the Catalog (2005 - 2016) are currently labeled with the category “Not reported” due to a lack of adequate information in the publication. Although confidentiality concerns certainly contribute to this, this large proportion of uncharacterized samples supports the notion that guidelines for the reporting of ancestry data are an absolute necessity. For this reason, we offer recommendations to increase standardization of ancestry reporting, with an emphasis on genetically-inferred ancestry, in publications (Box 1). We encourage implementation by authors reporting ancestry data and by editors reviewing publications that include human subjects.

## Conclusions

Genome-wide association studies have been enormously successful. However, the lack of clarity regarding the ancestry of samples and the lack of studies including diverse ancestral backgrounds raises questions about the interpretation and generalizability of results across populations. The framework we provide aims to address these challenges. It improves standardization of ancestry, increases integration of data, supports the assessment of diversity in large sets and facilitates further analyses. Its widespread adoption will enable the scientific community to investigate the generalizability of trait-associations across diverse populations, to identify associations unique to specific ancestries, to identify novel variants with clinical implications, and to help pinpoint causative variants, thus increasing our understanding of common diseases.

## Methods

### GWAS Catalog data curation

Details of GWAS publication identification, GWAS Catalog eligibility criteria and curation methods can be found on the GWAS Catalog website[40]. Extracted information encompasses publication information, study cohort information, including ancestry, and SNP-trait association results. Curation of ancestry data from the literature was performed according to Ancestry Extraction Guidelines outlined in the Supplementary Note.

### 1000 Genomes and HapMap Project population ancestry assignment

Information describing the 1000 Genomes[20] phase 3 and HapMap Project[19] phase 3 populations was taken from the Coriell Institute website[41]. Ancestry information, including ancestry category, country of recruitment, country of origin and additional information, was assigned to each population following the GWAS Catalog ancestry extraction guidelines (Supplementary Note).

### GWAS Catalog ancestry analysis

To determine the distribution of individuals, associations and traits by ancestry category, we first downloaded all Catalog data in tabular form[14]. All data (gwas-catalog-associations_ontology-annotated.tsv, gwas-catalog-ancestry.tsv, gwas-catalog-studies_ontology-associated.tsv, gwas-efo-trait-mappings.tsv) included in these analyses were curated from GWA studies published between 2005 and the end of 2015, with a release date of July 18 2017. The data can be found on the Catalog’s FTP site[42].

### Analysis of ancestry assessment methods in a subset of the GWAS Catalog

We selected the first 100 publications included in the Catalog (approximately covering the period between March 2005 to January 2008), and for comparison, the first 100 publications from 2016. For each publication, the method was assessed and classified into one of the following: 1. Self-reported, 2. Genetically assessed, 3. Ancestry stated without method, 4. Inferred from limited ancestry-related information (e.g. country information), 5. No ancestry information reported and 6. Mixed method (when a combination of methods was utilized to describe the study samples). Publications classified as “Genetically assessed” includes those where the author had clearly identified the genetic ancestry or admixture of the population, for example by using methods such as those described in Supplementary Box 1. It also includes those that confirmed self-reported information or defined samples based on self-reports but then excluded genetic outliers. Publications where no ancestry was stated, but curators inferred an ancestry based on country information are included in the fourth classification. In many cases authors used a statistical method to assess or control for ancestry or population stratification, without assigning individuals to a particular category, for example using a continuous axis of genetic variation from PCA to compute the association statistic. However, since this did not add any information that curators could use to assign a population ancestry to the study, it was not included under category 2.

## Declarations

### Ethics approval and consent to participate

Not Applicable

### Consent for publication

Not Applicable

### Availability of data and materials

The datasets generated and/or analyzed during the current study are available on the NHGRI-EBI GWAS Catalog search interface[4] and in spreadsheet form[14].

### Competing interests

PF is a member of the Scientific Advisory Board of Omicia, Inc.

## Funding

Research reported in this publication was supported by the National Human Genome Research Institute and the National Institute of General Medical Sciences of the National Institutes of Health under Award Numbers U41-HG007823 and U41-HG006104. The content is solely the responsibility of the authors and does not necessarily represent the official views of the National Institutes of Health. This research was also supported by the European Molecular Biology Laboratory. L.A.H., P.H. and H.J. are employees of the National Human Genome Research Institute.

## Authors’ contributions

J.M., J.A.L.M., P.H. H.A.J. and L.A.H. conceived this study and developed the ancestry framework. J.M., J.A.L.M., E.H.B., A.B., M.C., P.H., L.W.H., H.A.J., A.C.M., A.M. and L.A.H. performed curation of ancestry data of GWAS Catalog publications. J.M., J.A.L.M., M.C., T.B. and L.A.H. analyzed the distribution of ancestry categories in the Catalog and interpreted the data. L.W.H, J.A.L.M., L.H. and J.M. assessed the methods of ancestry determination utilized in GWAS Catalog studies and interpreted the data. A.C.M. and J.M. generated the figures. J.M., J.A.L.M and L.W.H. generated the Tables. E.H., D.W., C.M. and T.B. developed the GWAS Catalog curation and search interfaces. D.W. created the ancestry ontology, with contributions from J.M., J.A.L.M. and E.H.B. All authors contributed to the final manuscript, with J.M., J.A.L.M. and L.A.H. playing the key roles.

## Acknowledgements

The authors wish to thank all GWAS Catalog users and authors of studies included in the Catalog. We also thank Chris Gignoux for his expert review of the genomic methods of ancestry determination discussed in this manuscript and to Teri Manolio for valuable discussion.

## Acknowledgements

Research reported in this publication was supported by the National Human Genome Research Institute and the National Institute of General Medical Sciences of the National Institutes of Health under Award Numbers U41-HG007823 and U41-HG006104. The content is solely the responsibility of the authors and does not necessarily represent the official views of the National Institutes of Health. This research was also supported by the European Molecular Biology Laboratory. L.A.H., P.H. and H.J. are employees of the National Human Genome Research Institute. The authors wish to thank all GWAS Catalog users and authors of studies included in the Catalog. We also thank Chris Gignoux for his expert review of the genomic methods of ancestry determination discussed in this manuscript and to Teri Manolio for valuable discussion.

### Box 1. Recommendations for authors reporting ancestry data in publications

These recommendations were generated by expert curators following a detailed review of the over 3,000 GWAS publications included in the Catalog.

1. Preferentially use genomic methods to assess the ancestry of samples included in the GWAS Catalog. See Box 1 for a description of commonly used methods.
2. Indicate whether the background of participants was self-reported, inferred by genomic methods or a combination of both. If genetically inferred, indicate the analytical procedure utilized.
3. Provide detailed information for each distinct group of samples,

a. Ancestry descriptors should be as granular as possible (e.g. Yoruban instead of Sub-Saharan African, Japanese instead of Asian).
b. Avoid using country or citizenship as a substitute for ancestry.
c. Avoid using geographic descriptors that are part of a cohort name as a substitute for ancestry (e.g. TwinsUK cannot be assumed to be European ancestry).
d. If a population self-identifies using sociocultural descriptors (e.g. Old Order Amish), clearly state the genetic ancestry within which this sub-population falls.
e. If samples were derived from an isolated or founder population with limited genetic heterogeneity, clearly state the genetic ancestry within which this sub-population falls.
f. If available, genetic genealogy or ancestry of grandparents or parents should be included.
4. Assign an ancestry category for each distinct group of samples. See Table 1 for a list of ancestry categories. Refer to Supplementary Table 1 for a list of descriptors in use in the Catalog with their category assignments.
5. Provide the sample size for each distinct group of samples included in the analysis.
6. Provide country of recruitment.
7. If ancestry information is not available due to confidentiality, or any other concerns, note this in the publication.

## Figures

1. Figure 1 – Representation of ancestry data in the GWAS Catalog search interface
2. Figure 2 – Ancestry category distribution in the GWAS Catalog

a. Figure 2a - Distribution of individuals by ancestry category
b. Figure 2b - Distribution of studies by ancestry category
c. Figure 2c - Distribution of associations by ancestry category
3. Figure 3 – Distribution of individuals in GWAS Catalog studies published between 2005 – 2010 compared to 2011 – 2016.

